# Virtual meetings promise to eliminate the geographical and administrative barriers and increase accessibility, diversity, and inclusivity

**DOI:** 10.1101/2021.07.07.451408

**Authors:** Juncheng Wu, Anushka Rajesh, Yu-Ning Huang, Karishma Chhugani, Rajesh Acharya, Kerui Peng, Ruth D. Johnson, Andrada Fiscutean, Carla Daniela Robles-Espinoza, Francisco M. De La Vega, Riyue Bao, Serghei Mangul

## Abstract

The COVID-19 pandemic brought a new set of unprecedented challenges not only for healthcare, education, and everyday jobs but also in terms of academic conferences. In this study, we investigate the effect of the broad adoption of virtual platforms for academic conferences as a response to COVID-19 restrictions. We show that virtual platforms enable higher participation from underrepresented minority groups, increased inclusion, and broader geographic distribution. We also discuss emerging challenges associated with the virtual conference format resulting in a decreased engagement of social activities, limited possibilities of cross-fertilization between participants, and reduced peer-to-peer interactions. Lastly, we conclude that a novel comprehensive approach needs to be adopted by the conference organizers to ensure increased accessibility, diversity, and inclusivity of post-pandemic conferences. Our findings provide evidence favoring a hybrid format for future conferences, marrying the strength of both in-person and virtual platforms.

## Introduction

The world today is facing unprecedented challenges owing to the COVID-19 pandemic. One important restriction in the scientific setting is the lack of physical interpersonal interactions, which forms the core of academic conferences. This situation has adversely affected most sectors of daily life including the manner in which academia and industry operate. Despite this, it is essential to continue sharing scientific knowledge, and the research community was quick to adapt to COVID-19 restrictions with the majority of conferences effectively adopting an online delivery, virtual format. Online platforms provide a viable solution to the problem of sharing knowledge remotely and enable virtual connections between scientists, and sharing code, data, and comments through them has become easier^1^. Additionally, online conferences help alleviate an environmental challenge: Prior to the COVID-19 pandemic, there were open discussions about the substantial carbon footprint associated with conferences, attributed mainly to air travel related to in-person attendance and which contributes to human-induced climate change^2^. Having conferences online means that there will be less travel involved, thus reducing the carbon footprint^3^. The advancement of technology over the past decade and the ability to attend a virtual conference from any PC or laptop without the need for custom hardware^4^ also added to the appeal of a virtual setup.

With these benefits in mind, as well as the convenience of attending global conferences from any location, many members of the scientific community had been pushing for scientific meetings to be conducted at least in a hybrid manner - partly in-person and partly online - if not fully online^5^. However, this idea was not widely adopted then, perhaps due to the substantial logistics burden associated with ensuring proper internet access, organizing time zones, and an unwillingness to let go of the status quo. Nonetheless, out of necessity, conferences have now developed their own virtual platforms within a short time to accommodate the present challenges, often with support from the industry.

There are also logistic aspects to take into account. In some respects, more effort is required to put together a virtual conference than an in-person one, especially in terms of engaging prospective participants and garnering their interest for the upcoming years as well. However, while in-person conferences have a restriction on the number of attendees they can accommodate, a major advantage of virtual conferences is that the number of participants attending, and the geographical regions the conference can reach, can be scaled up. This advantage not only provides flexibility in who can attend but also promises to break logistical barriers associated with physical traveling and to connect researchers across the globe. Virtual conferences are not bound to one physical location, which promises to increase global participation and promote inclusivity^6,7^. Another major advantage is the reduction of cost, not only in terms of registration fees and travel by the attendees but also in terms of organizing the conference itself^8^. In this study, we set out to test this hypothesis by analyzing the demographics of attendees at four major conferences before and after the fully virtual format was adopted.

### Virtual platforms enable increased participation, diversity, and inclusion

What distinguishes a conference from a series of webinars is the active participation by the attendees of a conference^9^. We performed a systematic analysis of 24 conferences between January and August 2020 across medical, biology, computer science, and other fields. Out of these 24, 22 adapted to the virtual format, while two of the conferences were canceled altogether for the year. Among those 22, we observed a decrease in registration fee overall, with most (but not all) conferences waiving off the attendance fee altogether for the online conference. We selected the most popular conference in computational biology with free registration to illustrate the impact of virtual platforms on reducing financial and administrative burdens for participants. To further illustrate the distribution of participants and speakers across gender, ethnicity, and country comparing virtual with in-person platforms in previous years, we focused on four conferences, namely Bioinformatics Community Conference (BCC), BioConductor Annual Meeting (BioC), Intelligent Systems for Molecular Biology (ISMB), Research in Computational Molecular Biology (RECOMB) (**Figure 1, Table S1**).

**Figure 1.**
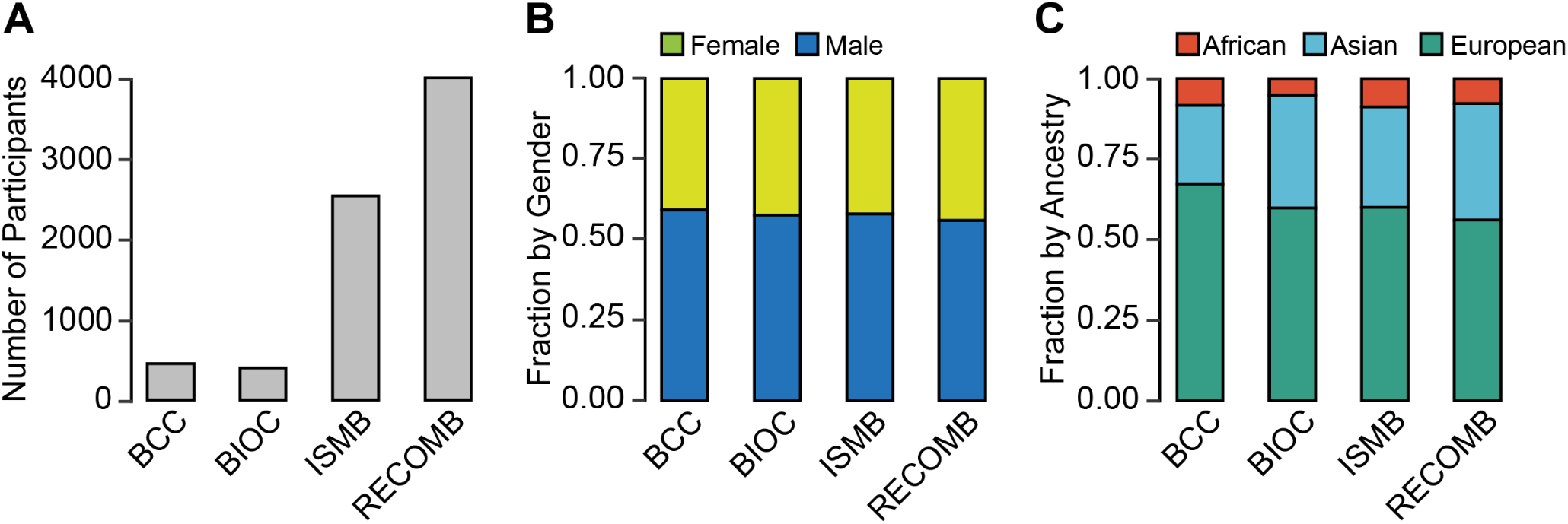
Distribution of participants by gender and ancestry for the 4 virtual-based conferences in 2020. (**A**) The total number of participants. (**B**) Percentage of females and males. (**C**) The number of participants by ancestry. BCC = Bioinformatics Community Conference. BIOC = BioConductor Annual Meeting. ISMB = Systems for Molecular Biology. RECOMB = Research in Computational Molecular Biology. African ancestry includes Africans, Muslim; Asian ancestry includes East Asian, Japanese, Indian Sub Continent; European ancestry includes British, East European, Jewish, French, Germanic, Hispanic, Italian, Nordic, as defined in the previous studies^15^.

To investigate the impact of virtual conferences in more depth, we compared the distribution of participants from the in-person vs virtual platforms for one of the four conferences (RECOMB) with data from 2019 and 2020. The gender and ethnicity of participants were imputed from the names of the participants using machine learning approaches (see **Methods**). The total number of participants increased from 374 in 2019 to 3913 in 2020, an 900% increase. The percentage of female participants remains similar comparing 2020 with 2019 (**Figure 2**). The number of individuals belonging to underrepresented minorities (African American and Latinos) increased from 19 to 331, demonstrating a substantial increase in the virtual platform reaching a broader range of underprivileged communities. However, the relative proportion of attendants from underrepresented minorities remains low (6.15% [23/374] in 2019 and 9.89% [387/3913] in 2020) (**Figure 2**), which may indicate that additional obstacles impede the participation of those groups in academia, that were not improved after removal of financial (e.g., free registration) and administrative (e.g., no traveling requirement) barriers.

**Figure 2.**
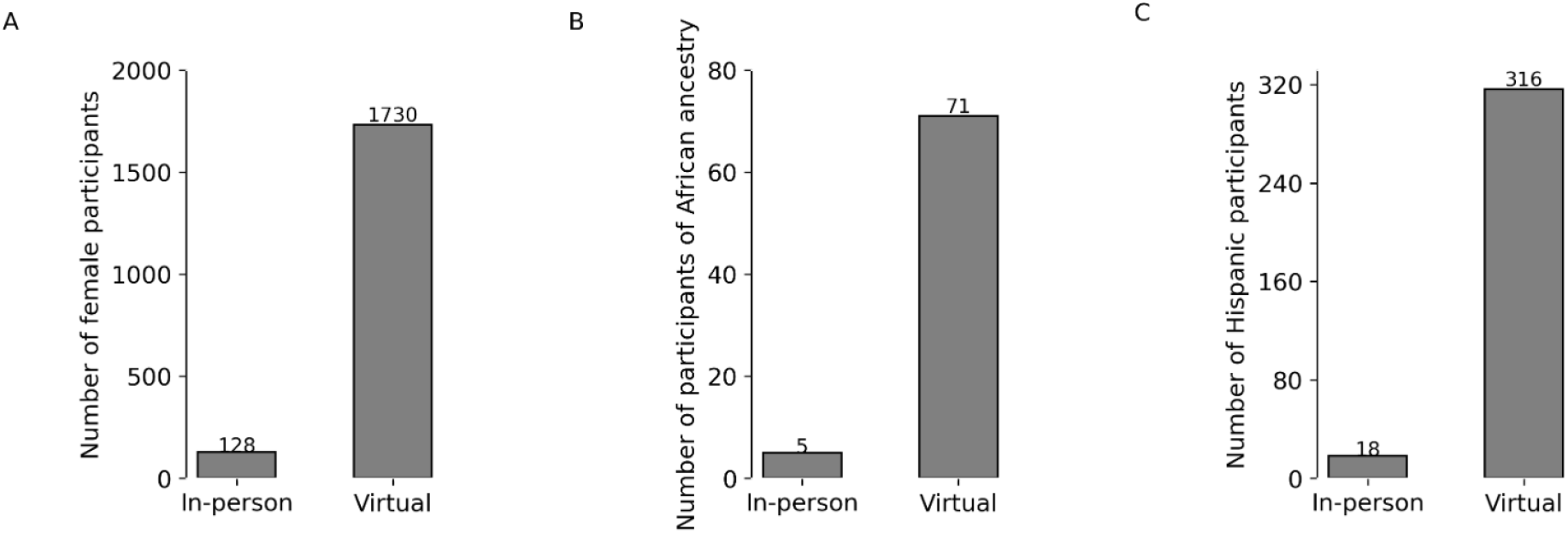
Underrepresented minority groups increase in 2020 (virtual) compared to 2019 (in-person) for the RECOMB conference. (**A**) Females. (**B**) Africans, (**C**) Hispanics. The number of participants is shown on the y-axis. Percentages are shown above each bar. Africans and Hispanics are subcategories of the African and European ancestry groups from **Figure 1**, respectively.

Accompanying the overall increase of participants, the geographical distribution has increased from 19 to 73 countries across the world, including a substantial increase in attendants from developing countries including those from Africa, South America, and Asia, including 3 low income and 35 middle-income countries (**Figure 3** and **4**). There is a substantial increase in the number of participants who joined the virtual conferences compared to the number of participants who joined the in-person conferences. (**Figure 4**) No participants from Oceania attended the in-person conferences, while 55 participants attended the virtual conferences. A similar trend for joining virtual or in-person conferences is also observed among participants from African countries. (**Figure 4, Figure S5**) Collectively, those results demonstrate an increased diversity due to virtual platforms which were not seen with in-person platforms.

**Figure 3.**
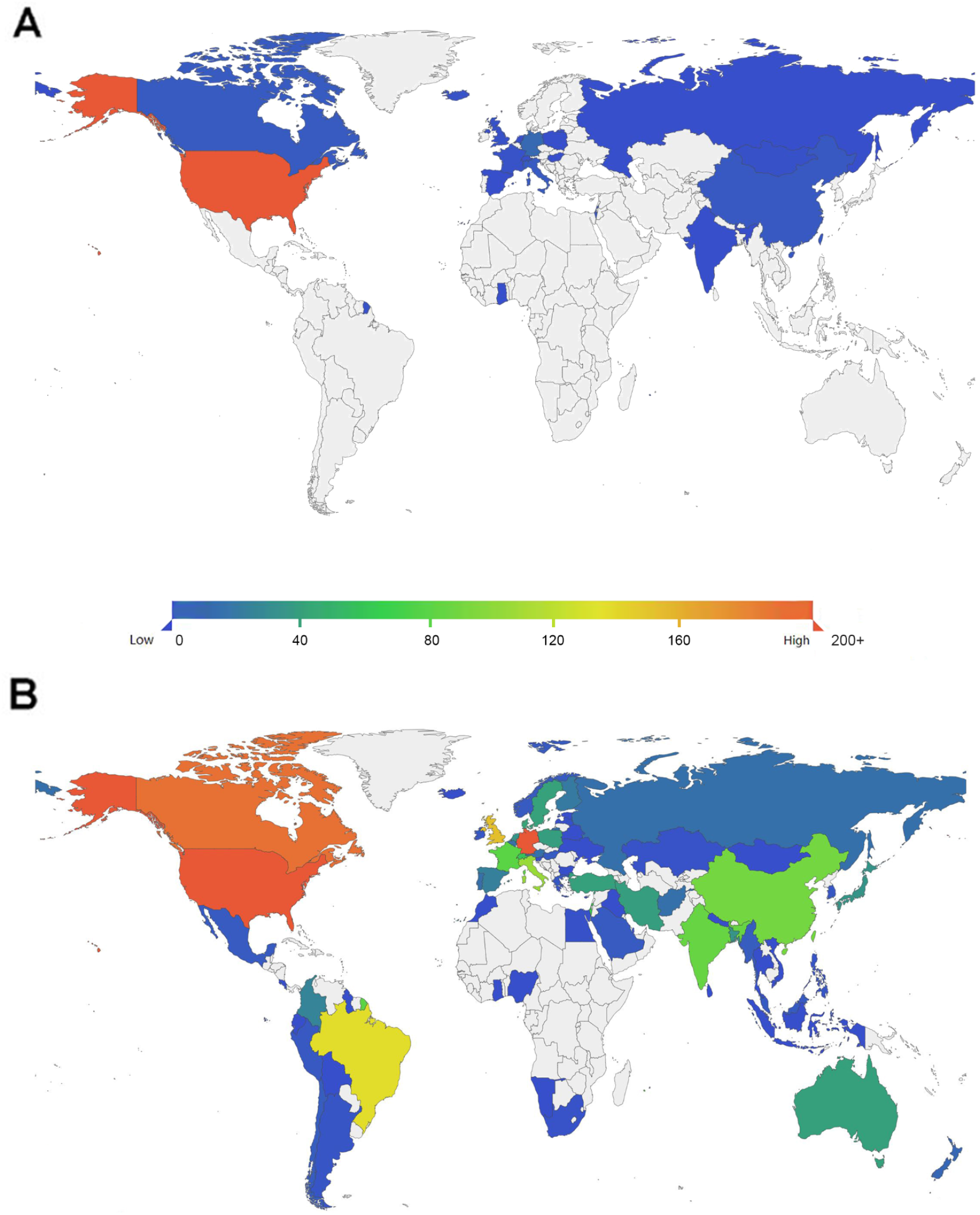
Increased diversity and inclusion of participants in the RECOMB conference in virtual compared to in-person formats. (**A** and **B**) Geographic distribution of the affiliations of participants in (**A**) 2019 and (**B**) 2020. Color represents the number of participants from each country (blue to red: 0 to over 200), as shown in the denotation bar at the left top corner in **A** and **B**.

**Figure 4.**
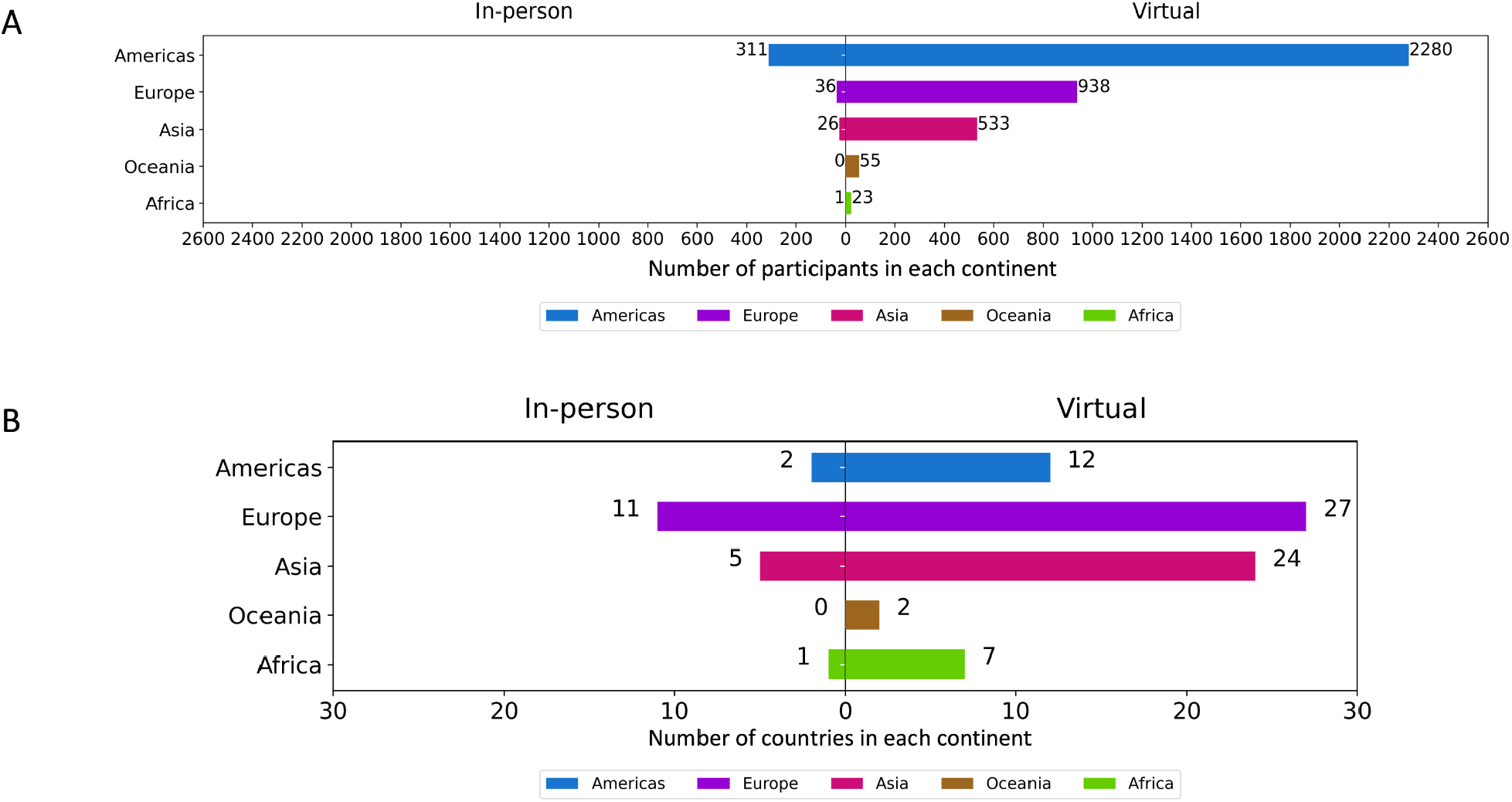
Comparison of the number of participants per country in virtual and in-person formats. (**A**) The absolute number of participants in each continent. (**B**) The absolute number of countries in each continent. (Color represents each continent, blue: Americas, purple: Europe, pink: Asia, brown: Oceania, green: Africa.) (Left: Participants that attended in-person conferences; Right: Participants that attended virtual conferences)

### Virtual platforms reduce the engagement of social activities and interactions

Inspired by the increased number of participants from virtual conferences, we explored the trend of online social activity as a proxy for measuring the engagement of participants in the conference activities. We hypothesized that more participants may lead to increased engagement in social media. However, this was not the case. Among three select large conferences of more than 1000 participants from computational biology, bioinformatics, and medical oncology (ASCO, AACR, ISMB, and BIOC), we quantified the number of tweets 3 days before, during, and 3 days after the dates of each conference for the past five to ten years (ASCO 2010-2020, **Figure S1**; AACR 2015-2021, **Figure S2**; ISMB 2015-2020, **Figure S3**; BIOC 2015-2020, **Figure S4**). For AACR, we collected the new data of 2021 as its conference had already occurred (April 2021) at the time of this study. We used hashtags to retrieve tweets relevant to each conference on Twitter (**Table S2**). We observed a continuing growth in related tweets from the earliest time until reaching the potential peak on the first or second day of the conference. During 2019 and 2020, we observed an overall decline of tweets in 2020 for ASCO, AACR, and ISMB (**Figures S1–S3**), while BIOC remains similar (**Figure S4**).

## Discussion

As a result of the ongoing pandemic, we have witnessed the evolution from a world where in-person meetings were the norm to a new reality in which nearly all conferences in the last year have been held virtually. There are positive aspects of this new format that support its continuation going forward. Online conferences increase participation from underrepresented scientists and those from developing countries and offer advantages such as schedule flexibility, reduced registration fees, and removal of travel barriers. Accordingly, in our analyses, we observed a dramatic increase in the number of participants from underrepresented minorities, as well as international attendants, including those from developing countries. The virtual conference platform offers increased accessibility and reduced administrative burden and more flexible programs compared to in-person ones, especially for international participants^7^. The remote format allowed more people to present their work and attend events by offering recorded lectures by on-demand online services and eliminating physical space limitations. All 22 conferences that we investigated provided recorded videos and workshop materials for an extended time after the conference ended, and even posted some of these materials to public media channels such as YouTube, open to the public permanently. In addition, select conferences such as the Keystone Symposia have published all recorded talks online free of charge to its members from developing countries, removing financial, administrative, and geographic barriers. An open science environment has also been promoted through Slack channels for conference reactions, conversations, and comments. The advantage of these online communication channels is that they are free, long-term, and sustained for as long as participants have the need.

One important feature of virtual conferences, which has not been available in in-person meetings, is live captions on screen, and CART and ASL interpretations. Captions in real-time provide higher accessibility to participants with disabilities and non-native English speakers and enhance participants’ attention and accurate understanding of the content covered by the speaker. However, we must note that live captioning could be an added expense and might not be an option if we were to keep the conferences affordable, and delayed crowd-sourcing has been considered as an option by some conferences to provide such service. In addition, including a communicator for the hearing impaired is recommended and has been implemented in select conferences and online seminars, improving accessibility for all participants. In addition, conferences often deposit recorded talks to online video platforms. An example here would be BioC, where automatic creation of captions is available via speech recognition technology. This is dramatically beneficial and should be considered for future conferences.

However, this change has not been without its difficulties. We have witnessed new challenges associated to the virtual conference platform, including requirements for high-speed internet, reduced peer-to-peer interactions, the need to spend substantial periods of time in front of a computer that causes ‘screen fatigue’^10^, work and home responsibilities, lack of social interactions^11^ that would otherwise be possible in an in-person setting, and difficulty to tune in to live sessions if on a different, inconvenient time zone. Disability, visa requirements, travel times and cost, and inflexible schedules are among the barriers that stop researchers from underrepresented groups and developing countries from participating in scientific and medical conferences internationally^12^. Further, there are two significant drawbacks of virtual conferences: First, the reduced opportunity for cross-fertilization between specialties, which is when participants stick to their sessions of interest and do not wander into other, perhaps unrelated sessions, as it happens in in-person meetings. Secondly, the limited networking opportunities of casual interactions in poster halls or social events, which can lead to new collaborations, awareness of new findings, and career advancement. This implies that in-person conferences would always have an advantage with respect to these aspects, and when we conceptualize the hybrid conference format going ahead, we need to ensure that they are maintained. Therefore, we believe that organizers and societies should continue striving to provide travel fellowships and services^7^ (such as child care) to allow broad access to the in-person experience.

While virtual conferences reduce costs for attendees, another aspect to consider is the cost/benefit for organizers. Academic conferences are typically organized by professional societies (e.g. AACR, ISCB, etc.) and these conferences contribute significantly to their revenue and membership, which in turn benefit the societies’ mission, programs, and initiatives such as lobbying for research funding, developing statements, and standards, promoting and delivering education to its members, organizing outreach events, and providing travel and other types of fellowships. On one hand, as compared to in-person events, virtual conferences result in cost savings in food and beverage costs, poster board costs, and rental fees. Audiovisual costs are however comparable between in-person and virtual events; virtual events add more logistics complexity and time from organizers, technicians, and presenters for rehearsals and capturing, uploading, and quality control of videos. A further downside for organizers is that virtual events may not drive memberships in the same way as in-person events. For example, ISCB memberships dropped in 2020 from the levels of 2019 and don’t appear to be on track for recovering in 2021. Furthermore, part of the costs and revenues from academic conferences are often covered by sponsorships from industry and other organizations. It has been difficult to convince sponsors of equal return of investment in virtual conferences, and when sponsors participate, their contributions are smaller. Combined with a reduction in registration fees, virtual conferences lead to reduced overall revenues for organizers as compared to in-person meetings.

The finding that social media engagement during the conferences did not increase as the number of participants did was surprising. However, we acknowledge that our method might not reflect all social media activity from those conferences given some also have their own communication platform outside of Twitter. One possible explanation for such a phenomenon is that unlike in-person conferences, some virtual conferences might use different ways for participants to communicate besides social media. We also acknowledge that our twitter data does not normalize for increasing usage over time. In addition, different time zones may present logistic challenges for participants from other countries. While we observed a higher number of participants in virtual conferences, those active in social media may be fewer than those in in-person conferences, potentially due to challenges such as that it could be hard to completely separate day to day work from participating in a virtual conference online, or family-related duties, which could prevent participants (especially female) from being fully engaged in the conference. Thus, it can be assumed that while the participation has increased drastically, this does not necessarily mean that all attendees engage with the conference to a maximum extent. Since there is the convenience of attending the conference from home, participants may only concentrate on specific talks that garner their interest and not all of the talks, unlike in-person conferences.

Our findings provide evidence favoring a hybrid format for future conferences, marrying the strength of both in-person and virtual platforms. This would broaden the reach to more communities and a higher number of countries. Going forward, we want to advocate a hybrid mode of organizing conferences. While we strongly believe that in-person conferences have their own benefits, and that no online communication tool can mimic the in-person experience completely^13^, we cannot neglect the multiple advantages that online conferences offer - in addition to providing opportunities to previously underrepresented groups to attend global conferences, this will contribute towards decarbonizing conference travel after the pandemic^3,14^. In fact, several conferences have started to implement such a hybrid mode. The Medical Image Computing and Computing Assisted Intervention Society (MICCAI), announced both in-person and virtual plans though had to cancel the former due to uncertainties of the pandemic. The Society for Immunotherapy of Cancer, The Society for Melanoma Research (SMR), the CSHL Genome Informatics, and AGBT Precision Medicine all announced hybrid formats for their meetings. These stand as evidence for the growing need and acceptance of the hybrid conference format. Taken together, our study warrants a continuation of evaluation on data from future conferences with different platforms (in-person, virtual, or hybrid) to evaluate its influence in accessibility, inclusion, and diversity.

## Methods

### Study Approval

This study was approved on 1/19/2021 by the University of Southern California (USC) Institutional Review Board (IRB) and is exempted from IRB review under the USC Human Research Protection Program Flexibility Policy.

### Data Collection

We manually collected data from the online conferences’ registration system and event management platform for the participants’ list, affiliation, and country of residence. This data was processed with Python 3, and saved as tidy data, including full names and affiliations. We also downloaded 296240 tweets by related hashtags of five conferences over the past ten years using GetOldTweets3, a Python 3 library collecting data from the JSON provider Twitter used on browser search. The tool uses the scroll loader, which allows users to read more new tweets when they scroll down on a twitter webpage. The content of these new tweets are provided by a JSON file, which is used by GetOldTweets3 to collect tweet data for any search query or username. In this study, we used conference organizers’ name (ASCO: @ASCO, BioC: @Bioconductor, etc.) and official hashtags of each conference was used (ASCO: #ASCO2020, #ASCO20; ISMB: #ISMB2020, #ISMB20; full list of hashtags provided in **Table S2**, along with the time periods of 3 days before to 3 days after these conferences proceed each year in the query (BioC:2020-07-22 to 2020-08-04, etc.).

### Data Analysis

Based on the basic tidy data, we predicted gender with first names, predicted ethnicity with last names; predicted country, institution with affiliations. We performed gender prediction using a naive Bayes classifier. We randomly selected 500 names as the training set and another 500 names as the testing set from 7579 unique names with gender labels based on the Katrowitz names corpus of Python 3 library, Natural Language Toolkit. The ethnicity is predicted using Python 3 library ethnicolr, which uses a Long short-term memory (LSTM) model, an artificial recurrent neural network (RNN) architecture, associated with the US census data, the Florida voting registration data, and the Wikipedia data, to predict ethnicity. The country of each participant was retrieved through two different strategies. We first detected institution names in the professional affiliation of participants, then matched these names to the “university-domains-list”, an open-source JSON format data set created and managed by Hipo, which contains 9700 institutions’ names and their located countries. Through the matching, we identified countries for the majorities of participants. For entries with missing countries for participants from organizations that are not included in the “university-domains-list”, we used manual search and curation on Google with the names of each participant’s professional affiliations.

### Data and Code Availability

The code and datasets in this study are available at https://github.com/HCC-data-sciences-pub/virtual-conference-analysis.

## Acknowledgment

We thank Dr. Max Alekseyev for his assistance in data inquiry. We thank Diane Kovats, Chief Executive Officer of ISCB, for responding to our queries.

## Supplementary Tables

**Table S1.**
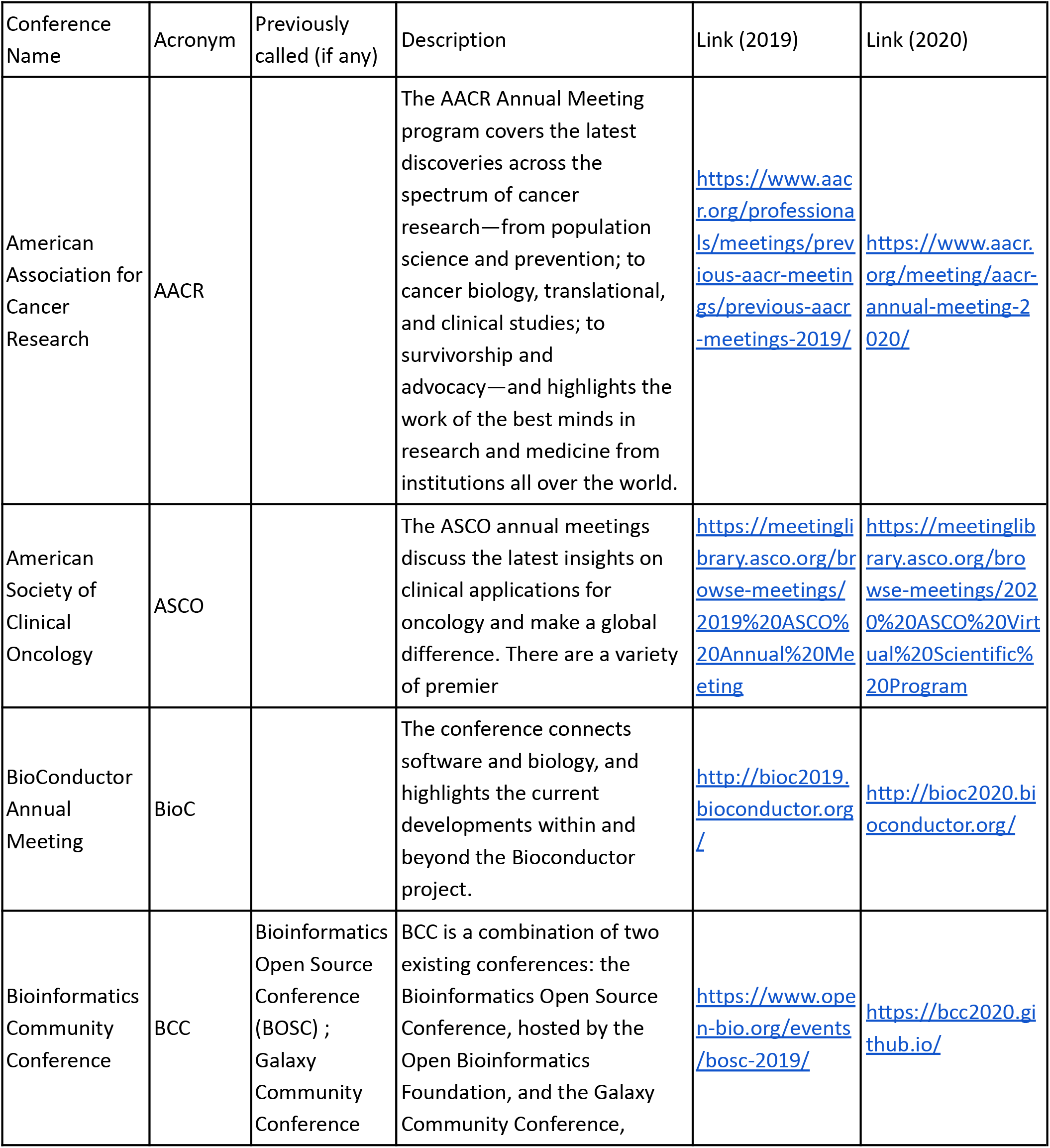
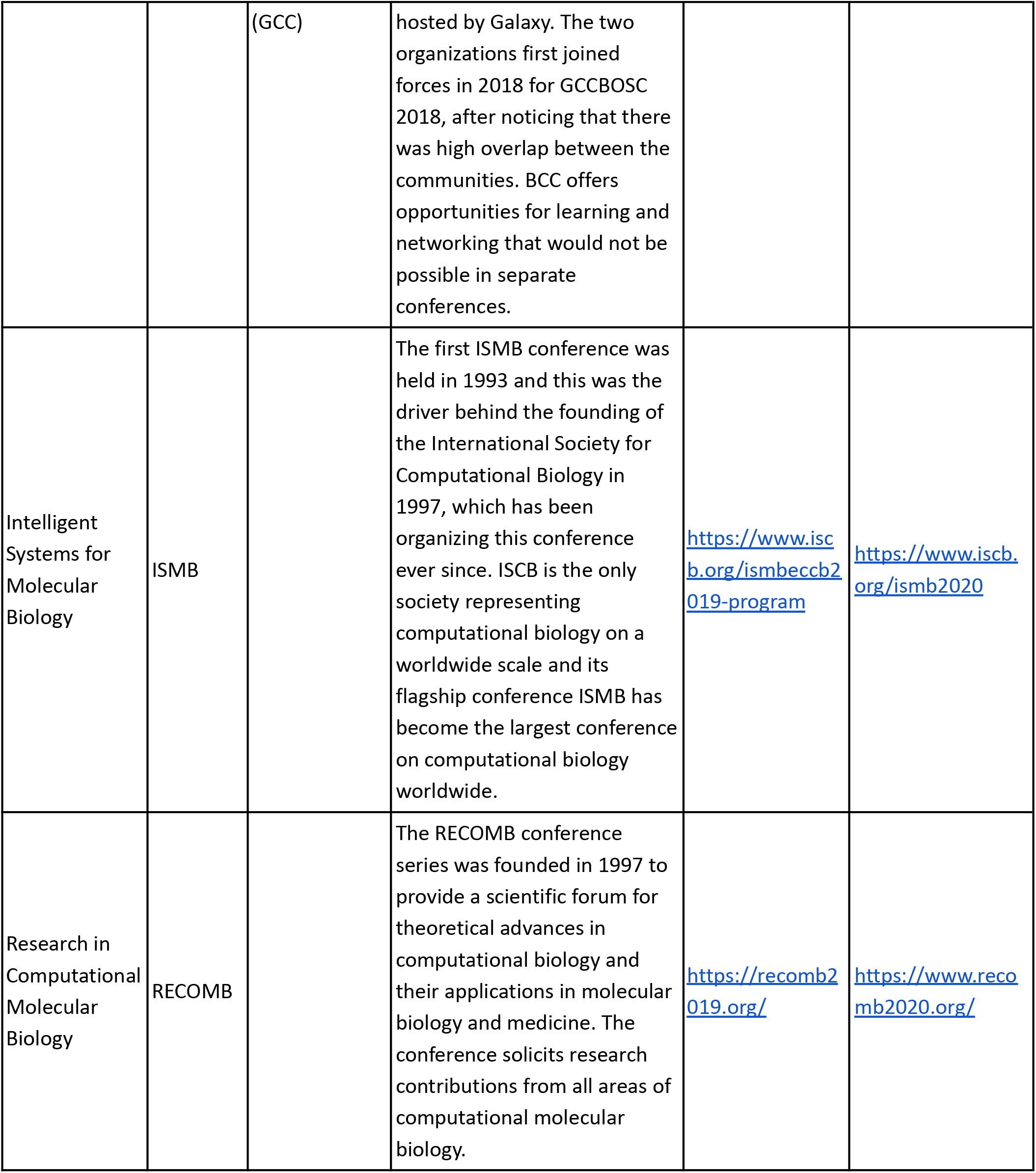
Description of the four conferences investigated in this study.

**Table S2.** List of hashtags used to retrieve tweets for conferences.

| Name | Search Query (i.e. 2020) | Year |
| --- | --- | --- |
| ASCO | "#ASCO2020 OR #ASCO20 OR @ASCO" | 2010-2020 |
| AACR | "#AACR2020 OR #AACR20 OR @AACR" | 2015-2021 |
| BioC | "#BioC2020 OR #BioC20 OR @Bioconductor" | 2014-2020 |
| ISMB | "#ISMB2020 OR #ISMB20 OR @iscb" | 2015- 2020 |

## Supplementary Figures

**Figure S1.**
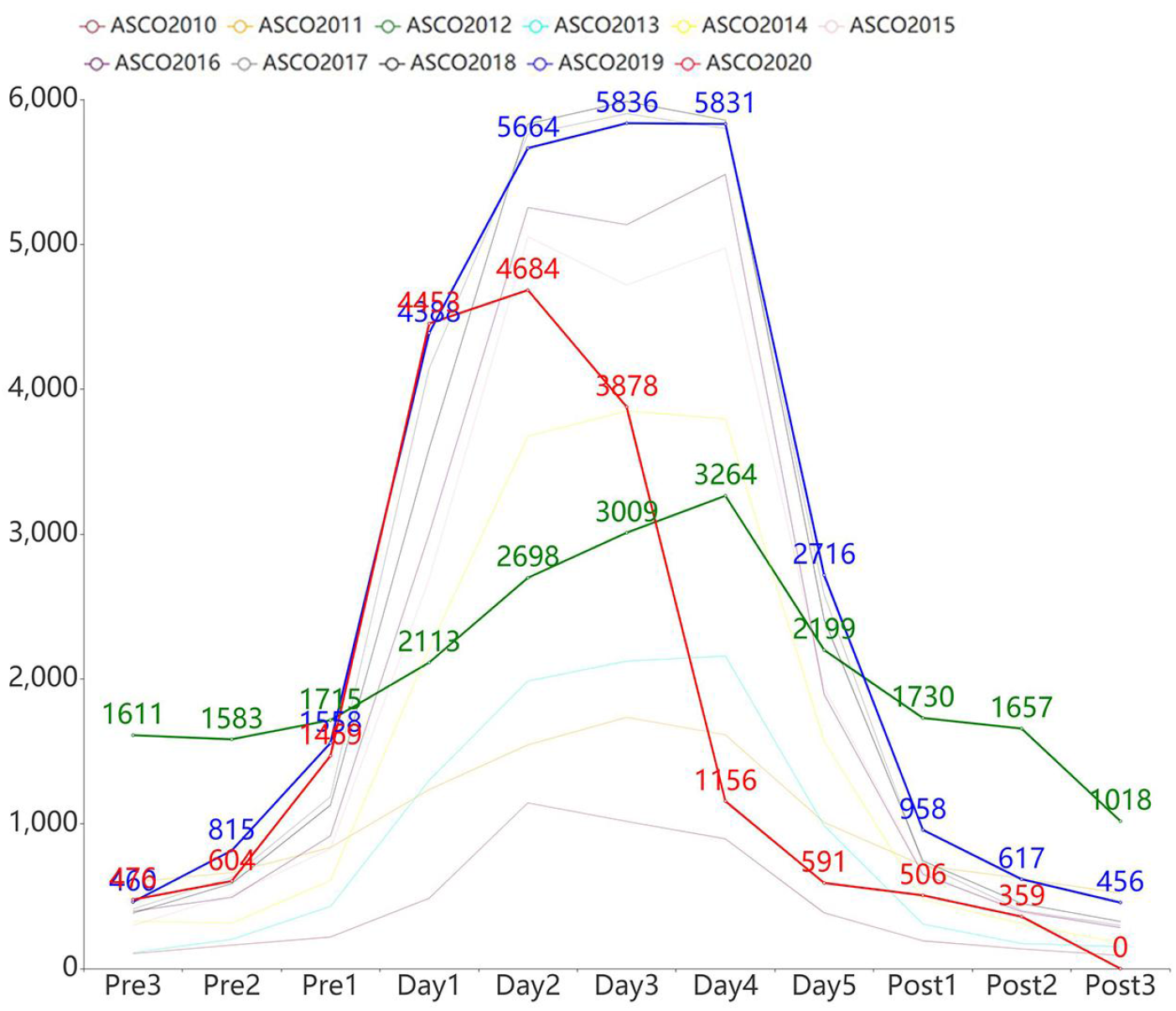
Number of tweets over time for the ASCO conference between 2010 and 2020. Three days pre-conference, five days during conference, and three days post-conference are shown. Each line represents one year. Three distinct trends are highlighted: The year 2020 (red), 2017 (blue), and 2012 (green).

**Figure S2.**
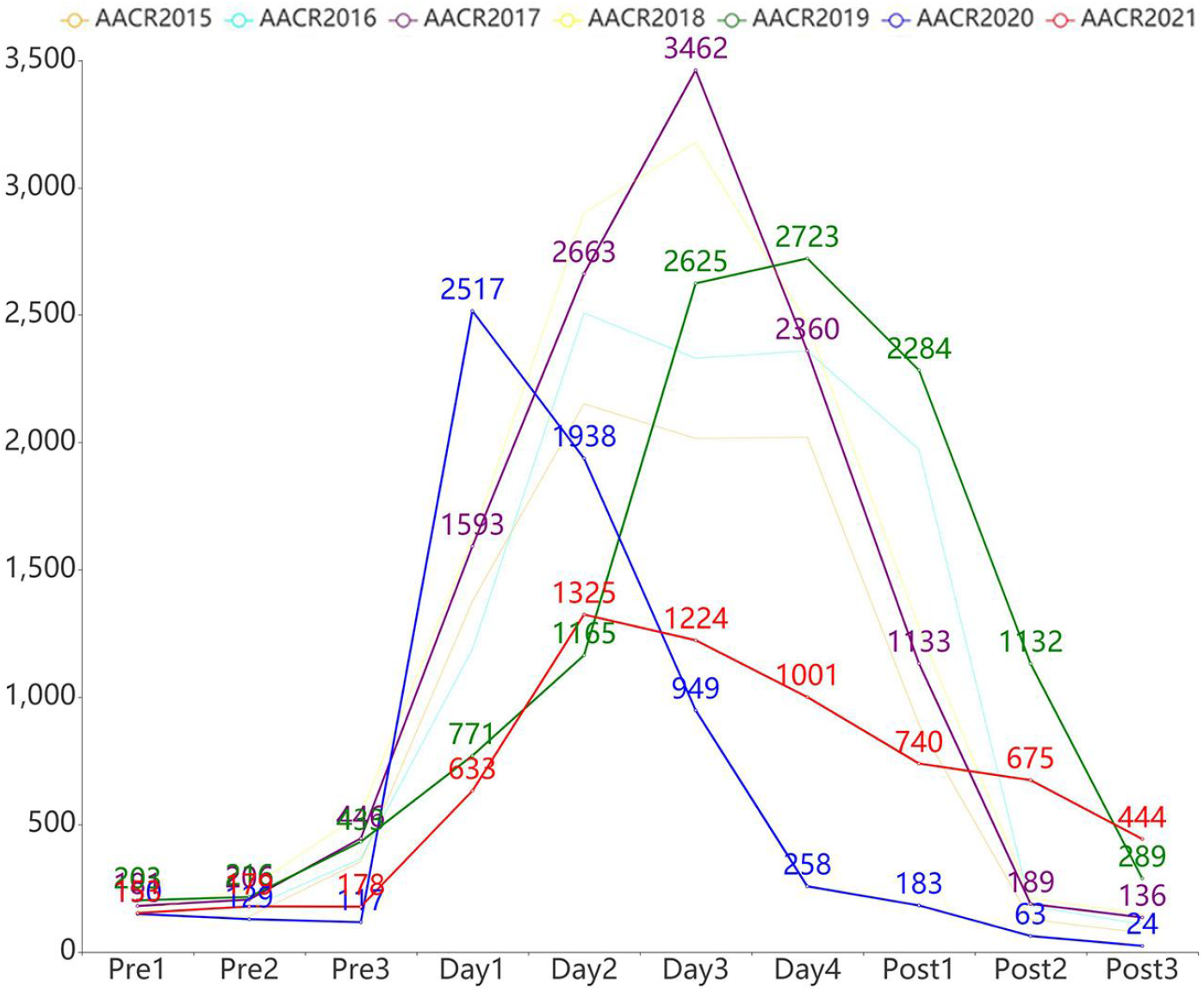
Number of tweets over time for the AACR conference between 2015 and 2021. Three days pre-conference, four days during conference, and three days post-conference are shown. Each line represents one year. Three distinct trends are highlighted: Year 2021 (red), 2020 (blue), 2019 (green), and 2017 (purple). Notice that for 2020 and 2021, the AACR Annual Meeting splits into two separate virtual meetings. We combined tweet numbers before, during, and after the conference time to align with other years.

**Figure S3.**
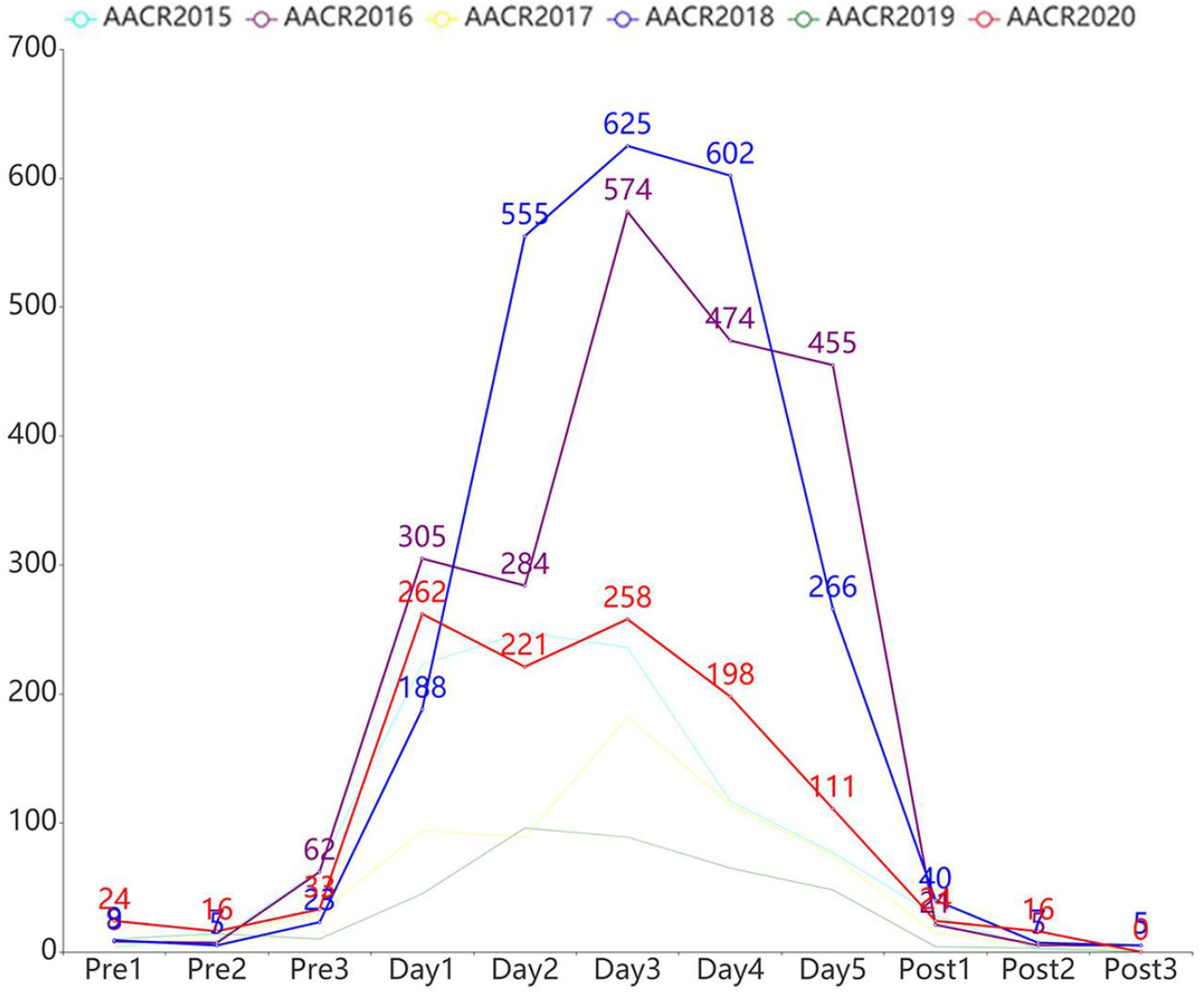
Number of tweets over time for the ISMB conference between 2015 and 2020. Three days pre-conference, five days during the conference (Besides 2020 which has only 4 days), and three days post-conference are shown. Each line represents one year. Three distinct trends are highlighted: The year 2020 (red), 2018 (blue), and 2016 (purple).

**Figure S4.**
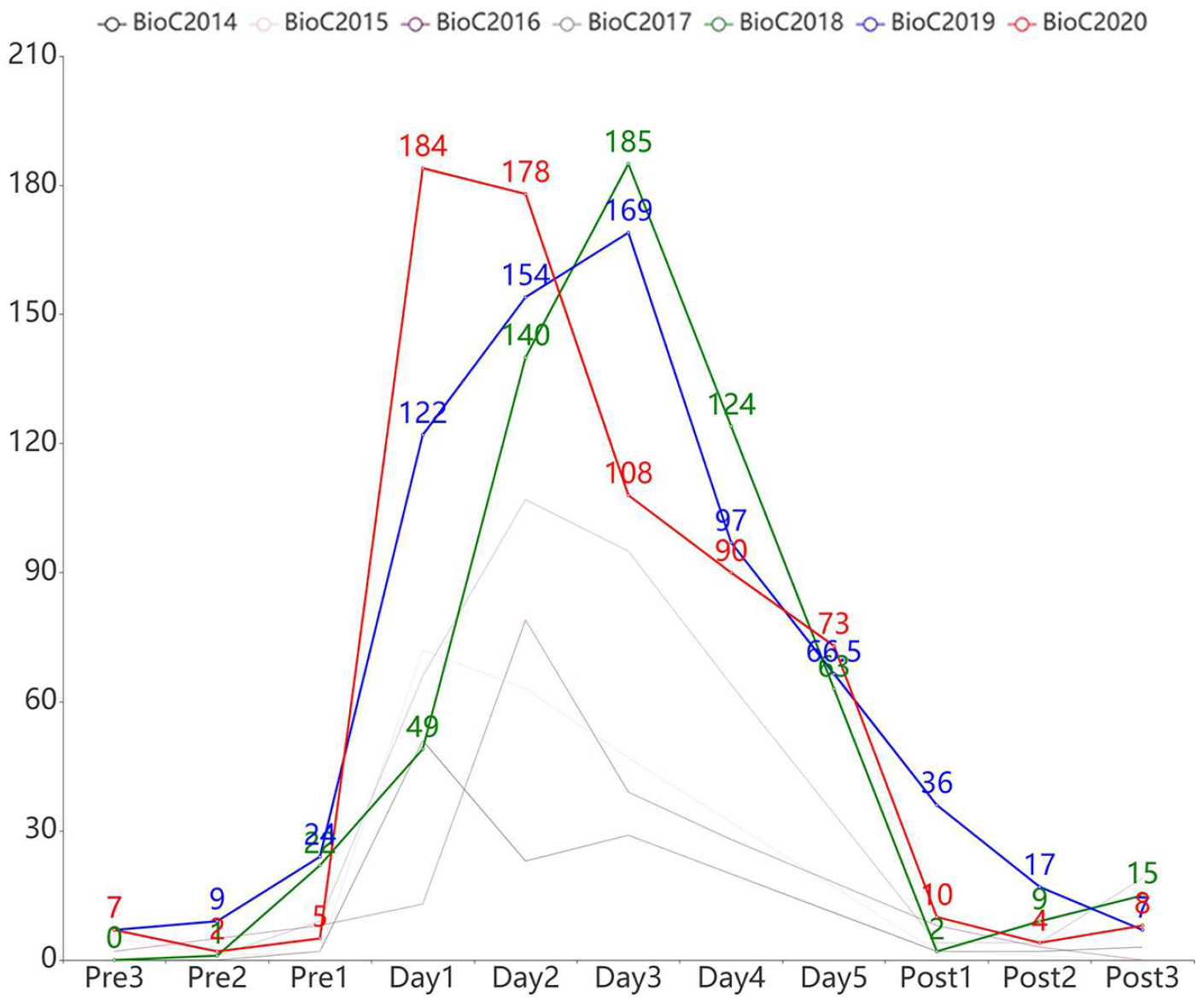
Number of tweets over time for the BioC conference between 2014 and 2020. Three days pre-conference, five days during the conference (2020 only, 4 days for 2019, 3 days for 2014-2018), and three days post-conference are shown. Each line represents one year. Three distinct trends are highlighted: The years 2020 (red), 2019 (blue), and 2018 (green).

**Figure S5.**
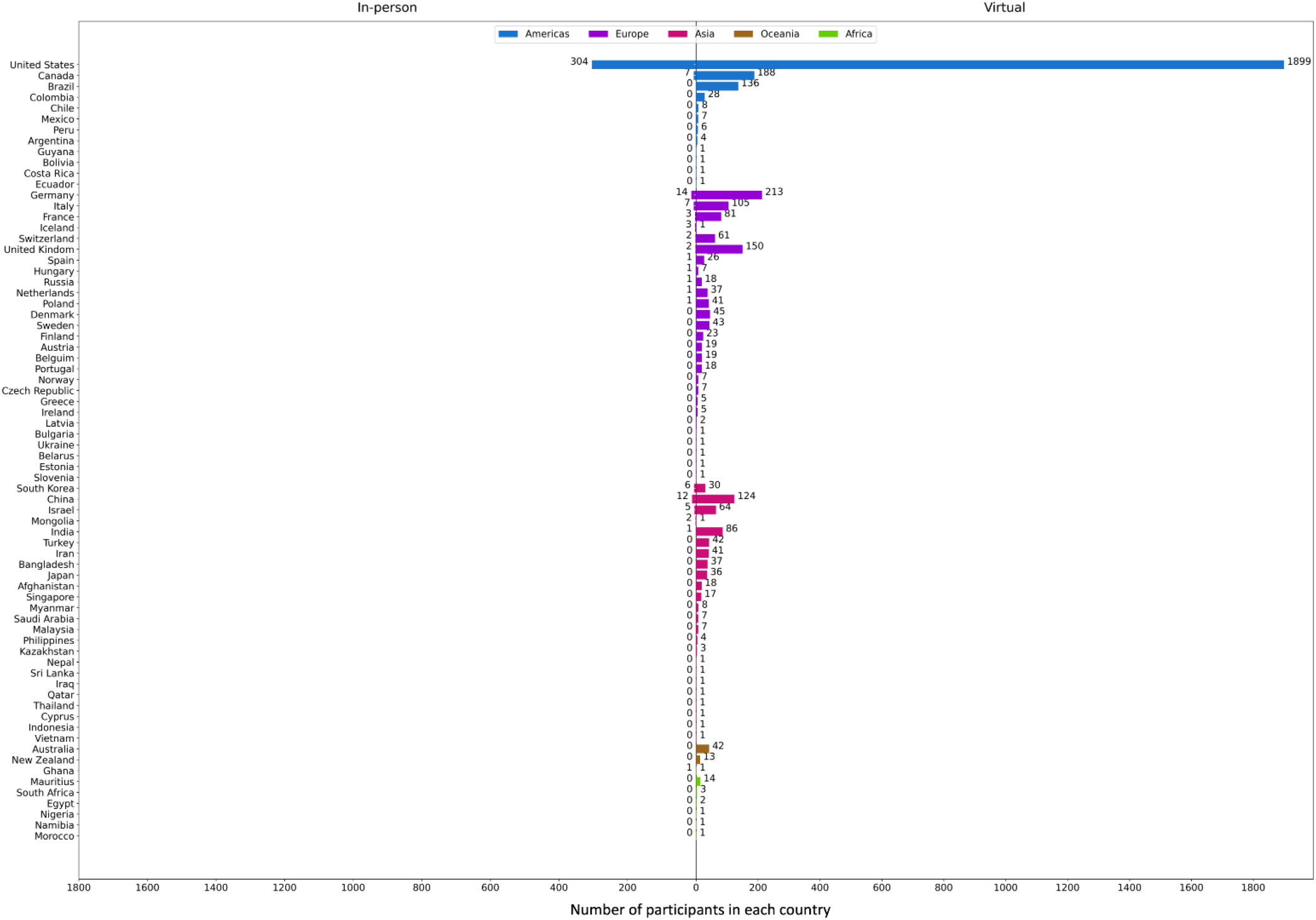
The absolute number of participants in each country. (Color represents each continent, blue: Americas, purple: Europe, pink: Asia, brown: Oceania, green: Africa.) (Left: Participants that attended in-person conferences; Right: Participants that attended virtual conferences)

## References

1. Sarabipour, S. Virtual conferences raise standards for accessibility and interactions. eLife 9, e62668 (2020).

2. Johnson, R., Fiscutean, A. & Mangul, S. Refining the conference experience for junior scientists in the wake of climate change. ArXiv200212268 Phys. (2020).

3. Barral, A. Virtual conferences are the future. Nat. Ecol. Evol. 4, 666–667 (2020).

4. Welch, C. J., Ray, S., Melendez, J., Fare, T. & Leach, M. Virtual conferences becoming a reality. Nat.Chem. 2, 148–152 (2010).

5. Newman, C. J. Post-COVID-19 scientific conferences: virtual becomes the new reality. Dev. Med.Child Neurol. 63, 493 (2021).

6. Levitis, E. et al. Centering inclusivity in the design of online conferences - An OHBM - Open Science perspective. (2021) doi:10.31234/osf.io/vj5tu.

7. Sarabipour, S. et al. Changing scientific meetings for the better. Nat. Hum. Behav. 5, 296–300 (2021).

8. Achakulvisut, T. et al. Towards Democratizing and Automating Online Conferences: Lessons from the Neuromatch Conferences. Trends Cogn. Sci. 25, 265–268 (2021).

9. Roos, G., Oláh, J., Ingle, R., Kobayashi, R. & Feldt, M. Online conferences – Towards a new (virtual)reality. Comput. Theor. Chem. 1189, 112975 (2020).

10. Foramitti, J., Drews, S., Klein, F. & Konc, T. The virtues of virtual conferences. J. Clean. Prod. 294, 126287 (2021).

11. Achakulvisut, T. et al. Improving on legacy conferences by moving online. eLife 9, e57892 (2020).

12. Pacchioni, G. Virtual conferences get real. Nat. Rev. Mater. 5, 167–168 (2020).

13. Ahn, S. J. (Grace) et al. IEEEVR2020: Exploring the First Steps Toward Standalone Virtual Conferences. Front. Virtual Real. 2, (2021).

14. Klöwer, M., Hopkins, D., Allen, M. & Higham, J. An analysis of ways to decarbonize conference travel after COVID-19. Nature 583, 356–359 (2020).

15. Ambekar, A., Ward, C., Mohammed, J., Male, S. & Skiena, S. Name-ethnicity classification from open sources. in Proceedings of the 15th ACM SIGKDD international conference on Knowledge discovery and data mining 49–58 (Association for Computing Machinery, 2009). doi:10.1145/1557019.1557032.

